## Supplementary for "Interrogating basal ganglia circuit function in Parkinson’s disease and dystonia"

**Supplementary Table 1: Data summary**

| **ID** | **disease** | **plasticity tasks** | **neurons** | **scale** | **total clinical score** | **hypokinetic clinical score** |
| --- | --- | --- | --- | --- | --- | --- |
| 1 | Parkinson’s disease | - | 5 | UPDRSIII | 34 | 14 |
| 2 | Parkinson’s disease | - | 3 | UPDRSIII | 32 | 10.5 |
| 3 | Parkinson’s disease | - | 7 | UPDRSIII | 48 | 17 |
| 4 | Parkinson’s disease | - | 6 | UPDRSIII | 39 | 16 |
| 5 | Parkinson’s disease | - | 1 | UPDRSIII | 40 | 19 |
| 6 | Parkinson’s disease | - | 7 | UPDRSIII | 47 | 23 |
| 7 | Parkinson’s disease | - | 5 | UPDRSIII | 45.5 | 17 |
| 8 | Parkinson’s disease | - | 10 | UPDRSIII | 31 | 14 |
| 9 | Parkinson’s disease | - | 12 | UPDRSIII | 34 | 11 |
| 10 | Parkinson’s disease | - | 9 | UPDRSIII | 29 | 11 |
| 11 | Parkinson’s disease | - | 7 | UPDRSIII | 26 | 10 |
| 12 | Parkinson’s disease | - | 3 | UPDRSIII | 46 | 21 |
| 13 | Parkinson’s disease | - | 4 | UPDRSIII | 37 | 18.5 |
| 14 | Parkinson’s disease | - | 2 | UPDRSIII | 48 | 17 |
| 15 | Parkinson’s disease | - | 5 | UPDRSIII | 29 | 8 |
| 16 | Parkinson’s disease | - | 2 | UPDRSIII | 68 | 27 |
| 17 | Parkinson’s disease | - | 3 | UPDRSIII | 31 | 11 |
| 18 | Parkinson’s disease | - | 4 | UPDRSIII | 17 | 5 |
| 19 | Parkinson’s disease | - | 1 | UPDRSIII | 31 | 12 |
| 20 | Parkinson’s disease | 1 | 3 | UPDRSIII | 23 | 12 |
| 21 | Parkinson’s disease | - | 6 | UPDRSIII | 35 | 15 |
| 22 | Parkinson’s disease | - | 8 | UPDRSIII | 32 | 10.5 |
| 23 | Parkinson’s disease | - | 1 | UPDRSIII | 13.5 | 3.5 |
| 24 | Parkinson’s disease | - | 5 | UPDRSIII | 43 | 15 |
| 25 | Parkinson’s disease | - | 3 | UPDRSIII | 31.5 | 10 |
| 26 | Parkinson’s disease | 2 | - | UPDRSIII | - | - |
| 27 | Parkinson’s disease | - | 3 | UPDRSIII | 65.5 | 32 |
| 28 | Parkinson’s disease | 1 | 2 | UPDRSIII | 38 | 16 |
| 29 | Parkinson’s disease | - | 1 | UPDRSIII | 32 | 13 |
| 30 | Parkinson’s disease | 3 | 8 | UPDRSIII | 40.5 | 19 |
| 31 | Parkinson’s disease | 1 | 9 | UPDRSIII | 24 | 11 |
| 32 | Parkinson’s disease | - | 14 | UPDRSIII | 23 | 7 |
| 33 | Parkinson’s disease | 1 | 6 | UPDRSIII | 60 | 22 |
| 34 | Parkinson’s disease | 1 | - | UPDRSIII | - | - |
| 35 | Parkinson’s disease | 1 | 3 | UPDRSIII | 30 | 7 |
| 36 | Parkinson’s disease | - | 4 | UPDRSIII | 29 | 11 |
| 37 | Parkinson’s disease | - | 3 | UPDRSIII | 46 | 19 |
| 38 | Parkinson’s disease | - | 6 | UPDRSIII | 26.5 | 9 |
| 39 | Parkinson’s disease | - | 9 | UPDRSIII | 37.5 | 18 |
| 40 | Parkinson’s disease | 2 | 3 | UPDRSIII | 49.5 | 23 |
| 41 | Parkinson’s disease | - | 3 | UPDRSIII | 57.5 | 24 |
| 42 | Parkinson’s disease | - | 4 | UPDRSIII | 59 | 20 |
| 43 | Parkinson’s disease | - | 4 | UPDRSIII | 35.5 | 12.5 |
| 44 | Parkinson’s disease | - | 4 | UPDRSIII | 68 | 26.5 |
| 45 | Parkinson’s disease | 1 | 8 | UPDRSIII | 41 | 16.5 |
| 46 | Parkinson’s disease | - | 6 | UPDRSIII | 32.5 | 15 |
| 47 | tardive dystonia | - | 2 | BFMDRS | 12 | n/a |
| 48 | cervical dystonia | 1 | 8 | TWSTRS | 22 | n/a |
| 49 | segmental dystonia | 2 | 7 | BFMDRS | 21 | n/a |
| 50 | hemidystonia | 1 | 0 | BFMDRS | 27 | n/a |
| 51 | generalized dystonia | - | 2 | BFMDRS | 11 | n/a |
| 52 | generalized dystonia | 1 | 0 | BFMDRS | 35 | n/a |
| 53 | generalized dystonia | 1 | 0 | BFMDRS | 27 | n/a |
| 54 | generalized dystonia | - | 4 | BFMDRS | 36 | n/a |
| 55 | craniocervical dystonia | - | 7 | TWSTRS | 22 | n/a |
| 56 | cranial dystonia | 3 | - | BFMDRS | 32 | n/a |
| 57 | cervical dystonia | - | 8 | TWSTRS | 29 | n/a |
| 58 | cervical dystonia | - | 5 | TWSTRS | 19 | n/a |
| 59 | cervical dystonia | - | 9 | TWSTRS | 3 | n/a |
| 60 | cervical dystonia | - | 18 | TWSTRS | 15 | n/a |
| 61 | cervical dystonia | - | 6 | TWSTRS | 4 | n/a |
| 62 | cervical dystonia | 1 | 10 | TWSTRS | 34 | n/a |
| 63 | cervical dystonia | - | 12 | TWSTRS | 13 | n/a |
| 64 | cervical dystonia | - | 10 | TWSTRS | 20 | n/a |
| 65 | cervical dystonia | - | 5 | TWSTRS | 35 | n/a |
| 66 | cervical dystonia | 2 | 6 | TWSTRS | 14 | n/a |
| 67 | cervical dystonia | - | 6 | TWSTRS | 8 | n/a |
| 68 | cervical dystonia | - | 8 | TWSTRS | 16 | n/a |
| 69 | axial dystonia | - | 3 | BFMDRS | 3 | n/a |

**Supplementary Table 2: Neuronal feature summary**

| **feature** | **dystonia** (based on 135 neurons from 19 patients) | **PD** (based on 222 neurons from 44 patients) |
| --- | --- | --- |
| firing rate (Hz) | 70.57 ± 19.39 | 83.08 ± 16.46 |
| burst index | 5.40 ± 1.42 | 4.21 ± 1.27 |
| coefficient of variation | 0.54 ± 0.09 | 0.44 ± 0.07 |
| theta (dB) | -10.88 ± 0.93 | -10.86 ± 1.68 |
| alpha (dB) | -12.48 ± 1.10 | -12.70 ± 1.27 |
| low beta (dB) | -12.85 ± 1.04 | -13.29 ± 0.95 |
| high beta (dB) | -12.51 ± 0.51 | -12.74 ± 1.33 |

**Supplementary Table 3: Multiple comparison via Bonferroni correction.**

| **test** | **independent variable** | **dependent variable (neuronal feature)** | **original p-value** | **corrected p-value** |
| --- | --- | --- | --- | --- |
| 1 | disease | firing rate | 0.0252 | 0.1746 |
| 2 | disease | burst index | 0.0024 | 0.0168 |
| 3 | disease | coefficient of variation | 7.46e-5 | 0.0005 |
| 4 | disease | theta power | 0.6265 | 1 |
| 5 | disease | alpha power | 0.3649 | 1 |
| 6 | disease | low beta power | 0.1176 | 0.8232 |
| 7 | disease | high beta power | 0.4585 | 1 |

**Supplementary Table 4: Multiple comparison via Benjamini-Hochberg false discovery rate**

| **test** | **independent variable (neuronal feature)** | **dependent variable (clinical score)** | **original p-value** | **corrected p-value** |
| --- | --- | --- | --- | --- |
| 1 | theta power | dystonia | 0.0242 | 0.17892 |
| 2 | low beta power | PD hypokinetic | 0.033 | 0.17892 |
| 3 | coefficient of variation | dystonia | 0.0364 | 0.17892 |
| 4 | firing rate | dystonia | 0.037 | 0.17892 |
| 5 | low beta power | PD total | 0.0426 | 0.17892 |
| 6 | high beta power | PD hypokinetic | 0.0698 | 0.225 |
| 7 | alpha power | PD total | 0.075 | 0.225 |
| 8 | alpha power | PD hypokinetic | 0.0902 | 0.236775 |
| 9 | high beta power | PD total | 0.111 | 0.259 |
| 10 | burst index | dystonia | 0.1748 | 0.36708 |
| 11 | alpha power | dystonia | 0.3898 | 0.744163636 |
| 12 | theta power | PD total | 0.5666 | 0.99155 |
| 13 | firing rate | PD total | 0.6666 | 1 |
| 14 | coefficient of variation | PD total | 0.6828 | 1 |
| 15 | coefficient of variation | PD hypokinetic | 0.757 | 1 |
| 16 | firing rate | PD hypokinetic | 0.832 | 1 |
| 17 | burst index | PD total | 0.903 | 1 |
| 18 | low beta power | dystonia | 0.9054 | 1 |
| 19 | burst index | PD hypokinetic | 0.9402 | 1 |
| 20 | theta power | PD hypokinetic | 0.9924 | 1 |
| 21 | high beta power | dystonia | 1 | 1 |

**Supplementary Fig. 1:**

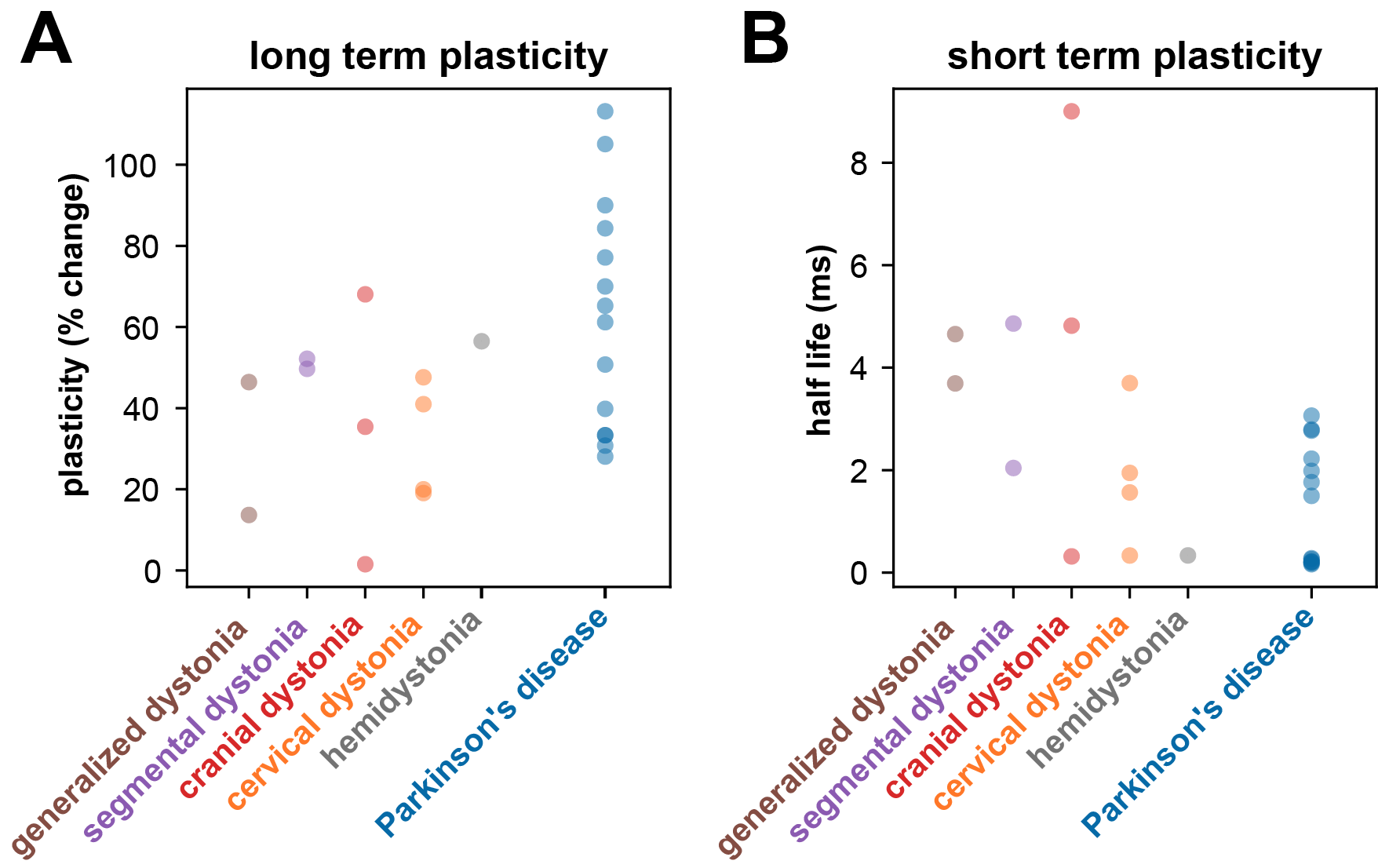

**Supplementary Fig. 1 –** **Long- and short-term plasticity across dystonia subtypes and PD.** (A) Shows the amount of plasticity (i.e., percentage change in fEP amplitudes pre- versus post-HFS) across different dystonia subtypes and PD. (B) Shows the half-life of the fitted exponential (i.e., rate of attenuation of fEP amplitudes) across different dystonia subtypes and PD.

**Supplementary Fig. 2:**

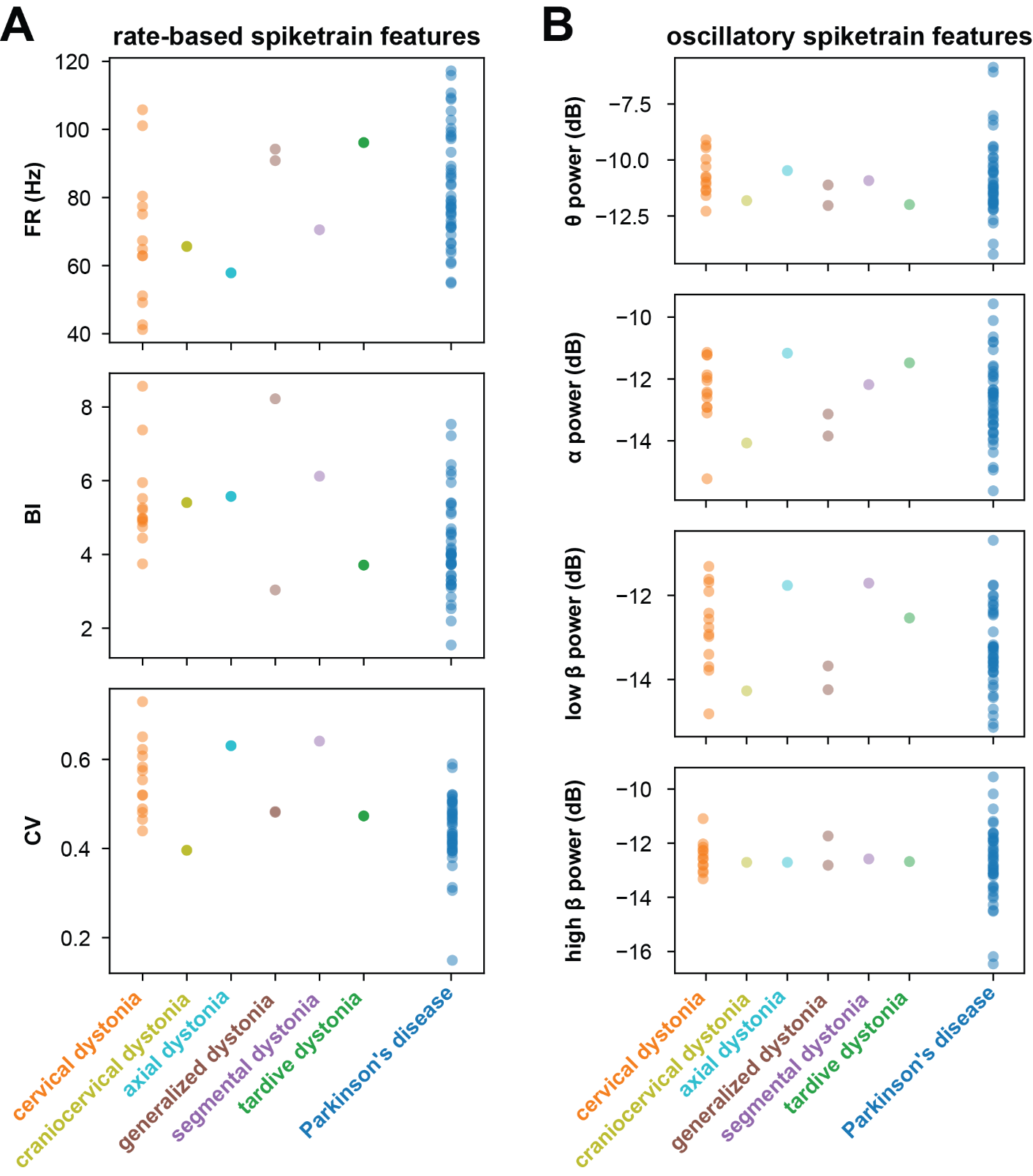

**Supplementary Fig. 2 – rate-based and oscillatory spiketrain features across dystonia subtypes and PD.** (A) displays rate-based features, including firing rate (FR), burst index (BI), and coefficient of variation (CV), across various dystonia subtypes and PD. (B) Illustrates oscillatory spiketrain features, such as theta, alpha, low beta, and high beta power, across different dystonia subtypes and PD.

**Supplementary Fig. 3:**

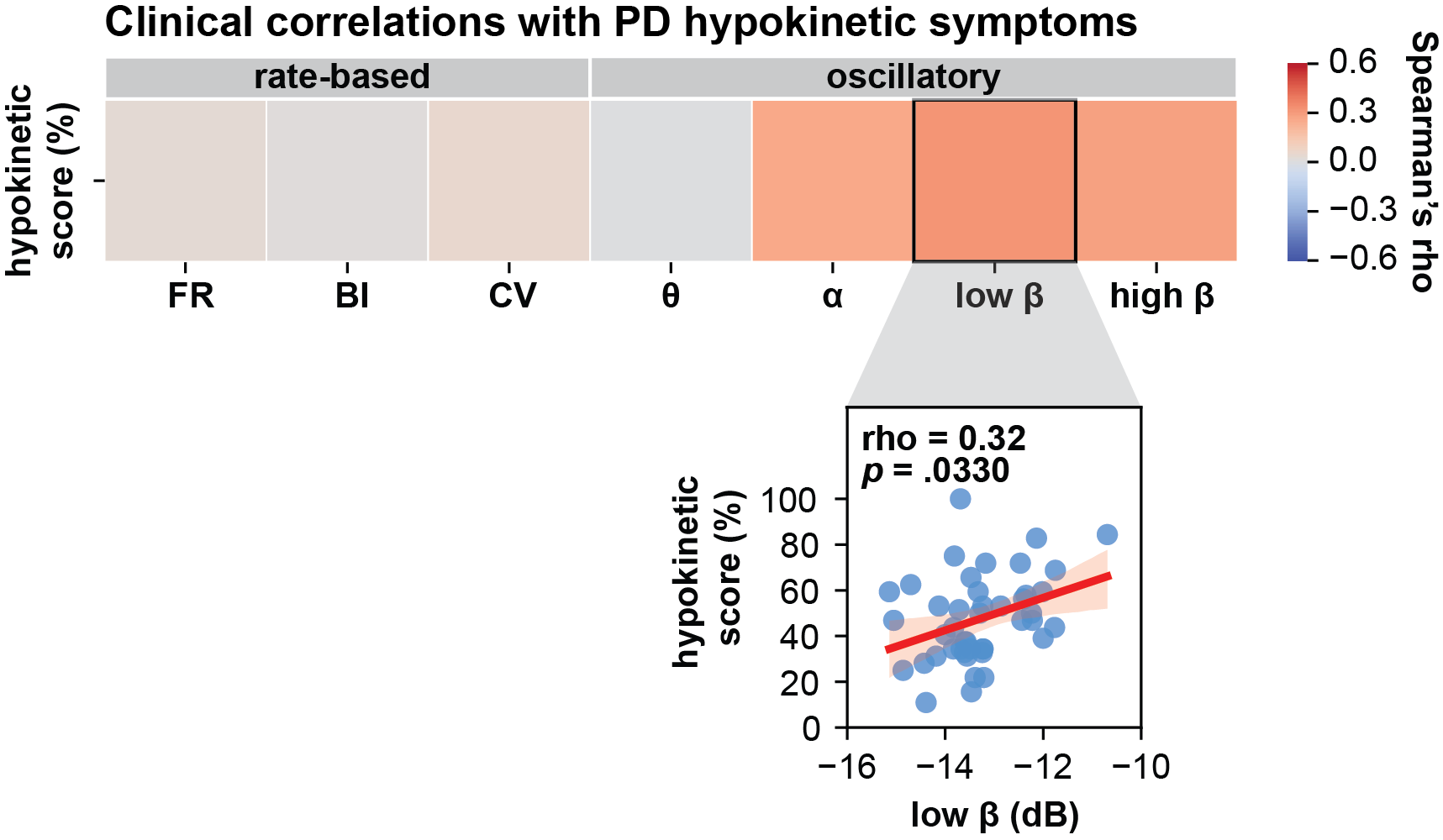

**Supplementary Fig. 3 – Clinical correlations with PD hypokinetic symptoms.** Low beta spiketrain oscillation power positively correlates with PD hypokinetic symptoms. The scatterplot depicts significant results before correction for multiple comparisons; however, this result did not withstand Benjamini-Hochberg correction for false discovery rate (refer to Supplementary Table 4).
